# Loss of SNORD115 mitigates SNORD116-driven sleep abnormalities in mouse models of Prader-Willi syndrome

**DOI:** 10.64898/2026.04.28.720143

**Authors:** Julie C. Riviere, Virginie Marty, Jérôme Cavaillé, Laure Verret

## Abstract

Prader-Willi syndrome (PWS) is a neurodevelopmental disorder caused by the loss of paternally expressed genes within the imprinted 15q11-q13 locus, which includes clusters of box C/D small nucleolar RNAs (SNORDs), notably the SNORD115 and SNORD116 gene families. Although paternally inherited SNORD116 deletions have been associated with sleep disturbances in patients and mouse models, the respective and combined contributions of SNORD115 and SNORD116 to sleep regulation remain unclear.

Here, we combined polysomnographic recordings with quantitative analysis of hypothalamic neuronal populations to assess sleep-wake architecture, sleep homeostasis, and underlying circuit alterations in mice carrying paternal deletions of SNORD115, SNORD116, or both clusters. SNORD116 deletion was associated with a selective increase in REM sleep, particularly during the light phase and during recovery following sleep deprivation, without affecting slow-wave sleep or REM-associated theta activity. In contrast, SNORD115 deletion did not alter sleep. Unexpectedly, combined deletion of SNORD115 and SNORD116 did not reproduce the REM sleep phenotype observed in SNORD116-deficient mice, indicating that SNORD115 loss attenuates SNORD116-dependent REM sleep alterations. At the cellular level, melanin-concentrating hormone (MCH) neuron density was reduced in both SNORD116-KO and double SNORD116/115-KO mice, whereas hypocretin (Hcrt) neurons were preserved across genotypes. Notably, REM sleep alterations did not parallel MCH loss, as increased REM sleep was absent in double-KO animals despite comparable reduction in MCH neuron density. Transcriptomic analyses at ZT0 further revealed only limited changes in hypothalamic gene expression across models.

Together, these findings reveal an unanticipated interaction between SNORD115 and SNORD116 in the regulation of REM sleep and uncover a dissociation between genetic alterations, neuronal circuit organization, and sleep phenotype. More broadly, they caution against inferring physiological functions or disease-relevant mechanisms from single SNORD deletions within complex imprinted loci, and indicate that current mouse models may not faithfully capture sleep alterations associated with PWS.

## Introduction

Prader-Willi syndrome (PWS) is a neurodevelopmental disorder initially characterized by perinatal growth retardation and hypotonia, followed by feeding disturbances (including hyperphagia), as well as hormonal and metabolic alterations that persist throughout life. This disorder results from the absence of expression of paternally inherited genes within the imprinted 15q11-q13 region, an epigenetically regulated chromatin domain containing multiple co-expressed protein-coding and noncoding RNA genes, including box C/D small nucleolar RNA (SNORD) genes (1,2) (**Figure 1**). Because several paternally expressed genes are simultaneously absent in most individuals with PWS, it remains unclear whether specific clinical features arise from the loss of a single critical gene or from the combined loss of multiple adjacent genes (3).

**Figure 1.**
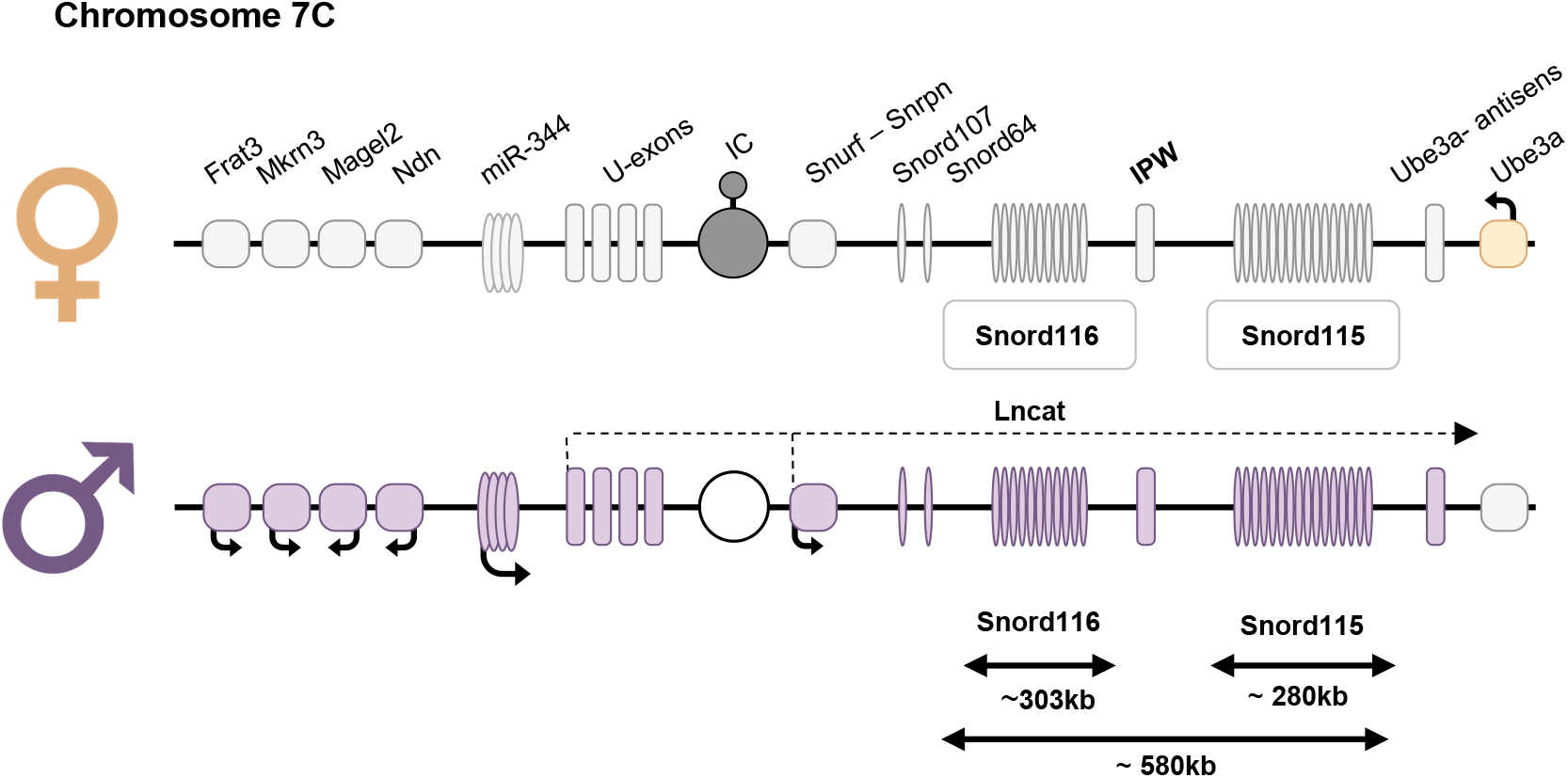
Schematic representation of the murine imprinted Snrpn domain. (chromosome 7), syntenic to the human Prader-Willi/Angelman syndrome (PWS/AS) locus (15q11-q13). This imprinted domain contains several protein-coding genes (square), long noncoding RNA genes (vertical rectangle), and small nucleolar RNAs (SNORDs; oval). SNORDs are embedded within introns of a long noncoding RNA (LNCAT). Paternal and maternal alleles are shown in purple and orange, respectively, with silent alleles in light gray. The imprinting center (IC) is unmethylated on the paternal chromosome and methylated on the maternal chromosome. The indicated deletions correspond to the SNORD116, SNORD115, and SNORD116/115 mouse models generated and used in this study. Not to scale.

The SNORD gene clusters, particularly SNORD115 and SNORD116, have received considerable attention. Rare individuals carrying microdeletions predominantly encompassing SNORD116 display core PWS features (4–10), positioning this cluster as a major contributor to the disorder. In contrast to most canonical SNORDs, which are highly conserved, ubiquitously broadly expressed and guide ribosomal RNA modification (ribose methylation, N4-acetylcytidine) through sequence complementarity (11–15), SNORD115 and SNORD116 are restricted to placental mammals, exhibit brain-specific expression in mice (16–18), and lack clearly identified RNA targets. As a result, their molecular functions and biological roles remain poorly understood in vivo (19–21). Current SNORD-deficient mouse models do not fully recapitulate the spectrum of PWS phenotypes, and studies in adult animals have largely focused on metabolic and feeding-related alterations (22–26; as reviewed in 27,28), leaving other domains, including sleep, comparatively unexplored.

Sleep disturbances represent one of the most disabling yet mechanistically unresolved dimensions of PWS (29–32). Excessive daytime sleepiness and shortened sleep onset latency are prominent features, which may persist despite treatment of obstructive sleep apnea (33–35). In parallel, alterations in REM sleep architecture, including REM fragmentation, have been reported in several case series (32,33,35,36). In some individuals, hypersomnolence is accompanied by cataplexy-like symptoms, suggesting a central disorder of arousal (36).

Many of these features have been linked to hypothalamic dysfunction (1). Sleep-wake regulation depends in part on two interacting neuronal populations in the lateral hypothalamic area (LHA): hypocretin/orexin (Hcrt) neurons, which promote wakefulness and food-seeking behavior (37–43) and melanin-concentrating hormone (MCH) neurons, which are directly involved in REM sleep regulation and contribute to feeding regulation (44–50). Reduced cerebrospinal fluid Hcrt levels have been reported in individuals with PWS and correlate with excessive daytime sleepiness severity (51,52), suggesting partial overlap with Hcrt-related sleep disorders such as type 1 narcolepsy (53). In parallel, MCH neurons have been implicated in the regulation of REM sleep, making these circuits particularly relevant to the alterations observed in PWS.

Consistent with a contribution of SNORD loss to sleep alterations, increased REM sleep duration and fragmentation, as well as elevated theta power during REM, have been reported in patients carrying microdeletions encompassing the SNORD116/115 interval (54). In parallel, studies using SNORD116-KO mouse models have described alterations in REM sleep architecture (55,56), whereas a role for SNORD115 in sleep regulation has yet to be demonstrated. Given that SNORD115 and SNORD116 are co-expressed in the brain and emerged together in placental mammals, they may exert overlapping or interacting functions. We therefore hypothesized that combined deletion of SNORD115 and SNORD116 might exacerbate sleep phenotypes observed in SNORD116-deficient mice. To test this, we directly compared mice carrying single deletions of either SNORD115 or SNORD116 with a novel model harboring a combined deletion of both clusters, hereafter referred to as SNORD116/115-KO (Marty et al., manuscript in preparation).

Unexpectedly, REM sleep alterations observed in SNORD116-deficient mice were not recapitulated in the double SNORD-KO model, revealing an unanticipated interaction between SNORD115 and SNORD116 loci. More broadly, these findings highlight the importance of studying multi-gene deletions in disease-relevant configurations such as PWS, particularly for sleep phenotypes where single-gene models may provide an incomplete or potentially misleading view of underlying mechanisms.

## Results

### SNORD116 deletion induces a limited increase in REM sleep

Previous studies have suggested that SNORD116 deficiency may contribute to sleep disturbances, including increased REM sleep and altered REM-associated theta activity in both PWS patients and SNORD116-KO mouse models (54,55). To assess the impact of SNORD116 deletion on sleep architecture, we performed 24-hour polysomnographic recordings in 2-month-old male SNORD116-KO mice and WT littermates (n = 8 per group) under baseline conditions.

Slow-wave sleep (SWS) architecture was unaffected by SNORD116 deletion. Total SWS duration, episode number, and delta power were comparable between genotypes, although strongly modulated by circadian phase (**Figure 2, A, B, C and D**). In contrast, REM sleep showed a small increase in SNORD116-KO mice. Total REM duration across the 24-hour cycle showed only a trend toward significance, but REM sleep was significantly elevated during the light phase (**Figure 2, E and F**), with KO animals spending approximately 8 minutes more in REM than WT littermates (8.9% vs. 7.9%). This difference was not accompanied by significant changes in REM episode number, mean episode duration (**Figure 2G and Supplemental Figure S1, A and D**), or theta power (**Figure 2H**), indicating that SNORD116 deletion induces a small increase in REM sleep amount without detectable changes in REM architecture.

**Figure 2.**
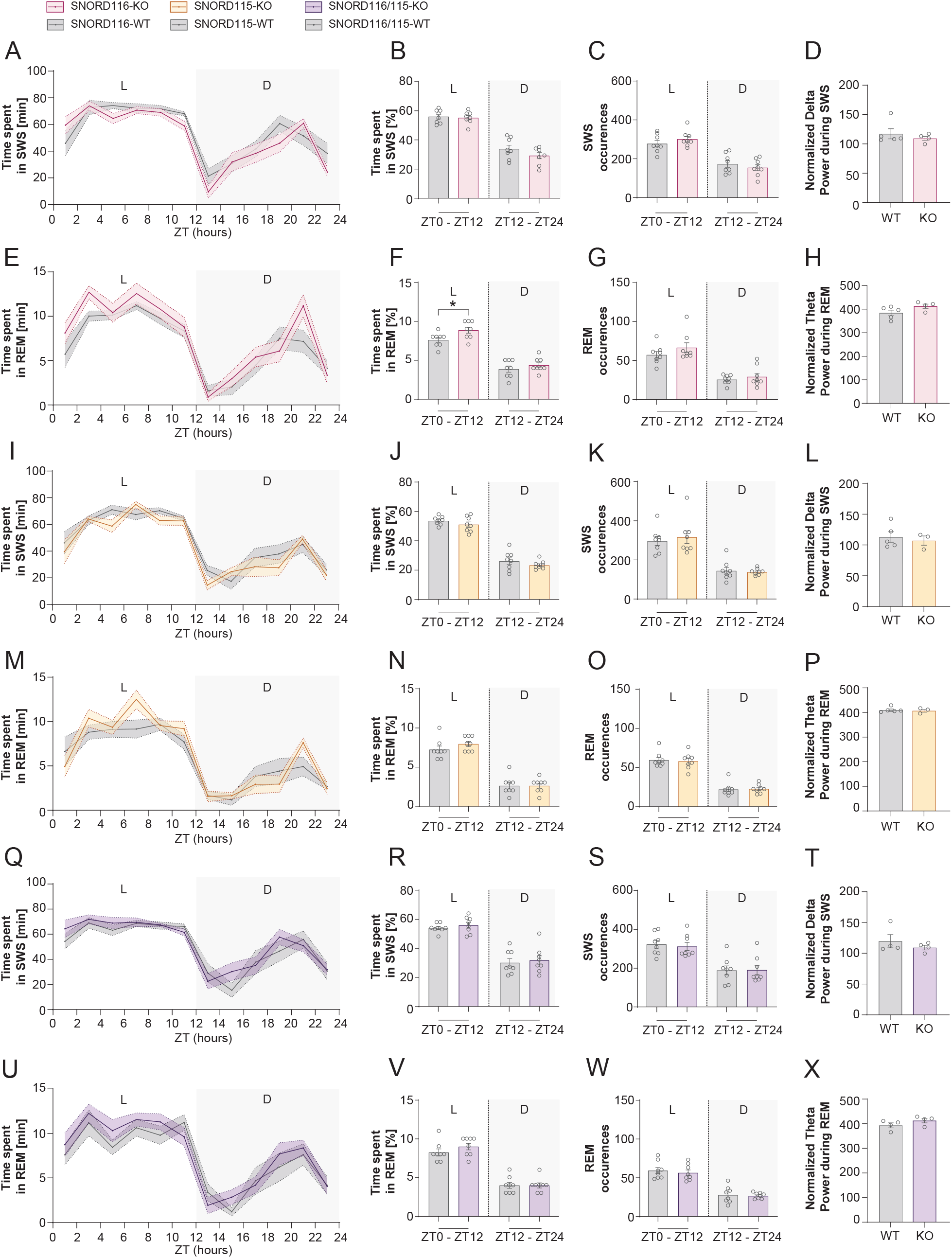
Vigilance state quantification, episode counts, and spectral analysis in SNORD116, SNORD115, and SNORD116/115 knockout mice under baseline conditions. Polysomnographic recordings were performed in 2-month-old male mice across three genotypes: SNORD116-KO, SNORD115-KO, and SNORD116/115 double-KO, with corresponding WT littermates. (**A-H**) **SNORD116 line:** (**A**) Time spent in slow-wave sleep (SWS; min) across the 24-hour cycle (2-hour bins, ZT0-ZT24). (**B**) Percentage of time spent in SWS during light and dark phases. (**C**) Number of SWS episodes. (**D**) Delta-band power (0.5-4 Hz) during SWS. (**E**) Total REM duration across 24 hours. (**F**) Percentage of time spent in REM sleep during light and dark phases. (**G**) Number of REM episodes. (**H**) Theta-band power (4-12 Hz) during REM sleep. (**I-P**) **SNORD115 line:** (**I-L**) SWS parameters as in A-D. (**M-P**) REM sleep parameters as in E-H. (**Q-X**) **SNORD116/115 double-KO line**: (**Q-T**) SWS parameters. (**U-X**) REM sleep parameters, including duration, episode number, and theta power. Data are presented as mean ± SEM. Time-course and light/dark phase data were analyzed using two-way repeated-measures ANOVA or Mixed-effects model (Genotype x Time or Phase) with Geisser-Greenhouse’s correction, followed by Šídák’s post hoc tests where appropriate. Spectral analyses were performed using unpaired t-tests with Welch’s correction or Mann-Whitney tests depending on data distribution. Sample sizes: n = 8 mice per genotype. *P < 0.05.

### SNORD115 deletion does not produce detectable changes in sleep architecture

We next assessed sleep architecture in 2-month-old SNORD115-KO mice and WT littermates. SNORD115 deletion had no detectable effect on sleep architecture. SWS parameters, including total duration, episode number, and delta power, were unchanged across genotypes (**Figure 2, I, J, K and L**). REM sleep was similarly unaffected, with no differences in total REM duration, its distribution across light and dark phases, REM episode number, or REM-associated theta power (**Figure 2, M, N, O and P**).

### Combined SNORD116/115 deletion does not reproduce the REM phenotype observed in SNORD116-KO mice

We next examined sleep architecture in SNORD116/115 double-KO mice and WT littermates (n = 8 per group). SWS parameters were unchanged, with no differences in total duration, episode number, or delta power compared to WT controls (**Figure 2, Q, R, S and T**). REM sleep was also comparable between genotypes. Total REM duration did not differ across the 24-hour cycle or between light and dark phases (**Figure 2, U and V**), and no differences were observed in REM episode number (**Figure 2W**) or REM-associated theta power (**Figure 2X**).

Thus, contrary to our initial hypothesis, the increase in REM sleep observed in SNORD116-KO mice was not detected in the double-KO model, despite the absence of a sleep phenotype in SNORD115-KO mice. This finding was confirmed in an independent cohort of older SNORD116/115-KO male mice (5 months; **Supplemental Figure S3**), supporting the robustness of this observation.

### SNORD116 deletion is associated with increased REM sleep during recovery following sleep deprivation

To assess sleep homeostasis, SNORD116-KO mice and WT littermates were subjected to a 4-hour sleep deprivation (SD) protocol from ZT0 to ZT4. SWS homeostasis was preserved in SNORD116-KO mice, with no differences in total SWS duration, episode number, or delta power compared to WT controls during the recovery period (ZT4-ZT10; **Figure 3, A, B, C and D**). In contrast, SNORD116-KO mice exhibited an increase in REM sleep during the recovery period compared to WT controls. During the first hours following SD (ZT4-ZT10), SNORD116-KO mice spent more time in REM sleep (~10-15 minutes), resulting in a higher amount of REM sleep during the light phase (**Figure 3, E and F**). This increase occurred in the absence of detectable changes in SWS homeostasis, indicating a preferential increase in REM during recovery. It was not associated with significant changes in REM episode number, mean episode duration (**Figure 3G and Supplemental Figure S2, A and C**), or REM-associated theta power (**Figure 3H**).

**Figure 3.**
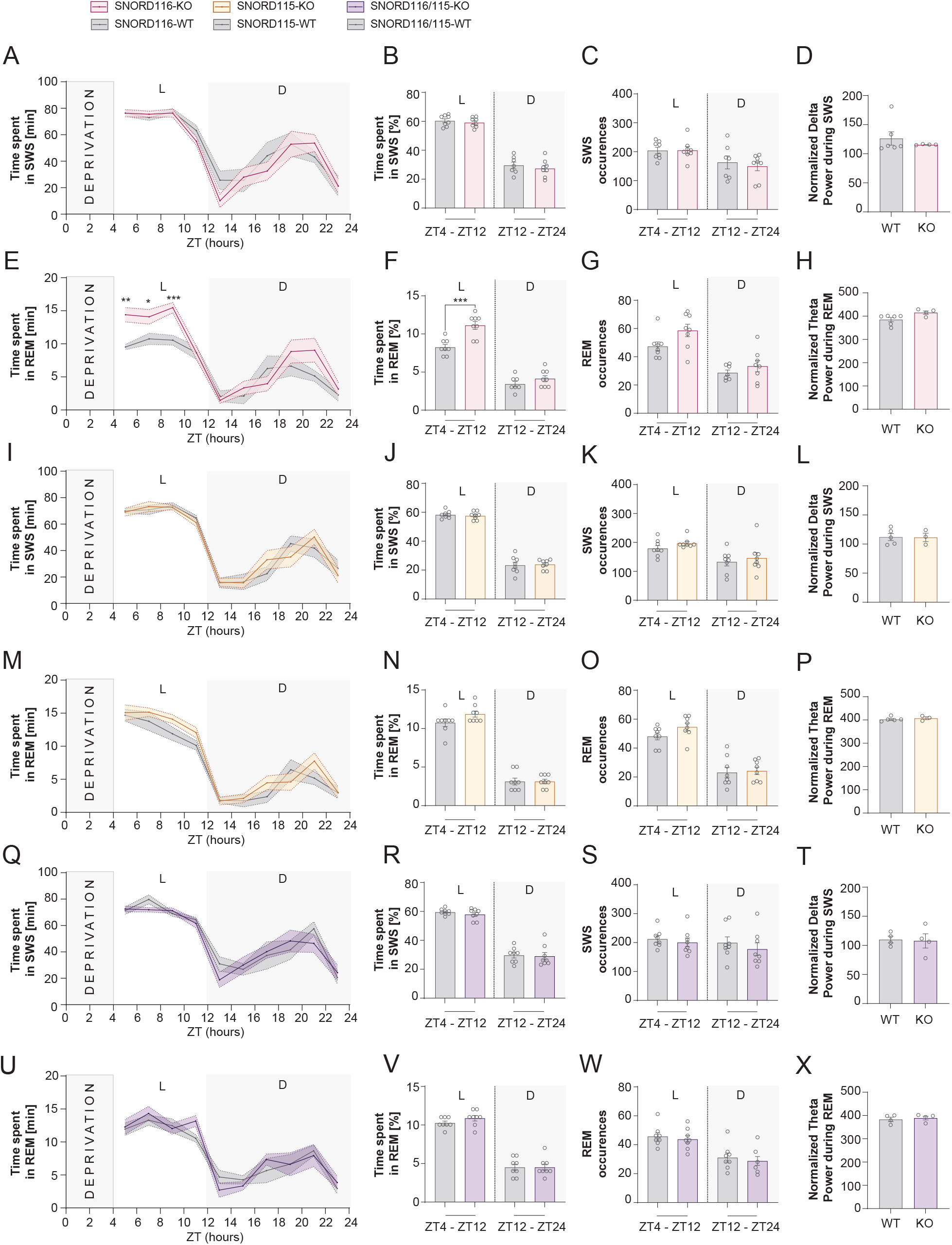
Sleep homeostasis following sleep deprivation in SNORD116, SNORD115, and SNORD116/115 knockout mice. Sleep homeostasis was assessed following 4 h of sleep deprivation (ZT0–ZT4) in 2-month-old male mice from SNORD116-KO, SNORD115-KO, and SNORD116/115 double-KO lines and their WT littermates. (**A–H**) **SNORD116 line: (A)** Time spent in SWS (min) across the recovery period (ZT4–ZT24; 2-hour bins). (**B**) Percentage of S WS during light (L) and dark (D) phases. (**C**) Number of SWS episodes across L/D phases. (**D**) Delta-band power (0.5–4 Hz) during SWS. (**E**) REM sleep duration across ZT bins. (**F**) Percentage of REM sleep during L/D phases. (**G**) Number of REM episodes. (**H**) Theta-band power (4–12 Hz) during REM sleep. (**I–P**) **SNORD115 line:** (**I–L**) SWS parameters as in A-D. (**M–P**) REM sleep parameters as in E-H. (**Q–X**) **SNORD116/115 double-KO line**: (**Q–T**) SWS parameters; (**U–X**) REM sleep parameters. Data are presented as mean ± SEM. Time-course (ZT bins) and L/D phase data were analyzed using two-way repeated-measures ANOVA (Genotype x Time or Phase), followed by appropriate post hoc tests. Spectral analyses were performed using unpaired t-tests with Welch’s correction or Mann–Whitney tests depending on data distribution. *P < 0.05; **P < 0.01; ***P<0.001.

### SNORD115 deletion does not alter REM or SWS homeostatic responses

SWS homeostasis was preserved in SNORD115-KO mice, with no differences in total SWS duration, episode number, or delta power compared to WT controls during the recovery period (ZT4-ZT10) (**Figure 3, I, J, K and L**). REM sleep was similarly unaffected, with no differences in REM duration during recovery, its distribution across light and dark phases, REM episode number, or REM-associated theta power between genotypes (**Figure 3, M, N, O and P**). In an independent cohort of older SNORD115-KO mice (7-8 months), SWS episode duration was modestly increased during the rebound period (**Supplemental Figure S4**), while other parameters remained unchanged.

### Combined SNORD116/115 deletion does not alter sleep homeostasis following sleep deprivation

SWS homeostasis was preserved in SNORD116/115 double-KO mice, with no differences in total SWS duration, episode number, or delta power compared to WT controls during the recovery period (**Figure 3, Q, R, S and T**). REM sleep was also unaffected following sleep deprivation. REM duration during recovery, its distribution across light and dark phases, REM episode number, and REM-associated theta power were all comparable between SNORD116/115 double-KO mice and WT littermates (**Figure 3, U, V, W and X**).

Unexpectedly, the increase in REM sleep observed in SNORD116-KO mice during recovery was not detected in the double-KO model.

### SNORD116 deletion reduces MCH neuron density in LHA without altering Hcrt neurons

To investigate whether sleep alterations were associated with changes in hypothalamic circuits, we quantified Hcrt and MCH neurons in the LHA across the three SNORD-deficient mouse lines.

Using combined immunohistochemistry and fluorescent *in situ* hybridization (FISH), we first confirmed that SNORD115 and SNORD116 are expressed in both Hcrt- and MCH-immunoreactive neurons (**Figure 4A**). Fluorescent signals appeared as punctate nuclear labeling with DAPI-poor regions, consistent with nucleolar localization (18,57). Signal specificity was verified using SNORD116/115 double-KO mice, which showed no detectable FISH signal (**Supplemental Figure S5;** 58).

**Figure 4.**
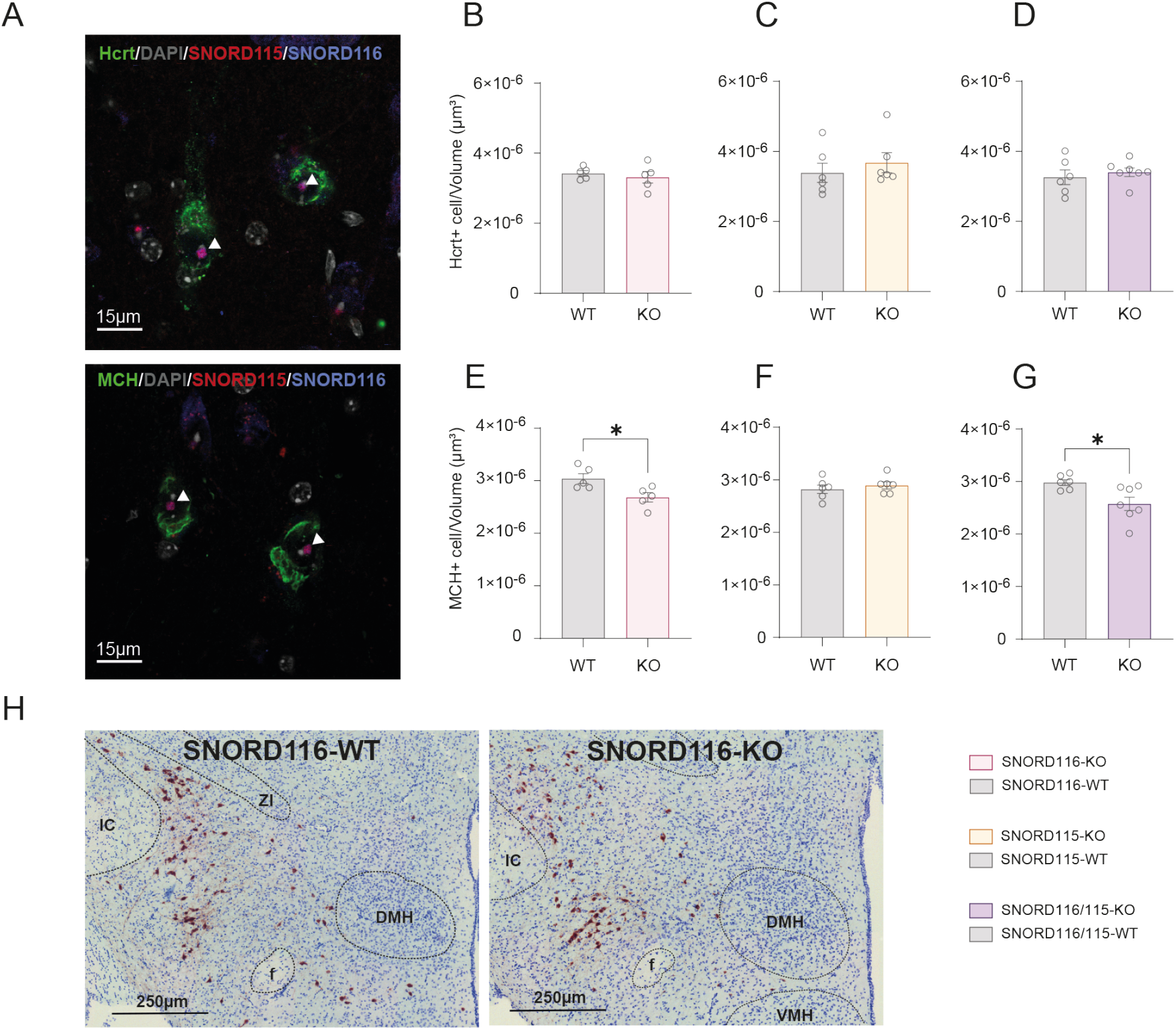
SNORD expression in Hcrt and MCH neurons and impact of SNORD loss on these hypothalamic neuronal populations. (**A**) Representative RNA FISH images showing SNORD115 and SNORD116 expression at the single nucleus level in brain sections from adult WT mice. SNORD115 and SNORD116 were detected using Cy3-(red) and Cy5-(blue)-labelled antisense DNA oligonucleotide probes, respectively, while hypocretin (Hcrt; top) and melanin-concentrating hormone (MCH; bottom) neurons were visualized by immunofluorescence (green). White arrows indicate the nucleolar compartments where SNORD115 and SNORD116 accumulate. (**B-D**) Quantification of Hcrt-positive (Hcrt^+^) neuron density within the lateral hypothalamic area (LHA; cells/µm^3^) in SNORD116-KO (**B**), SNORD115-KO (**D**), and SNORD116/115-KO (**E**) mice; no significant differences between genotypes. (**E-G**) Quantification of MCH-positive (MCH^+^) neuron density within the LHA (cells/µm^3^) in SNORD116-KO (**E**), SNORD115-KO (**F**) and in SNORD116/115-KO mice (**G**); a significant reduction was observed in SNORD116-KO and SNORD116/115-KO mice. (**H**) Representative coronal hypothalamic sections from SNORD116-WT (left) and SNORD116-KO (right) mice following anti-MCH immunostaining and cresyl violet counterstaining. Data are presented as mean ± SEM. Statistical analyses were performed using unpaired t-tests or Mann–Whitney tests depending on data distribution. *P < 0.05. Arc, arcuate nucleus; DMH, dorsomedial hypothalamic nucleus; IC, internal capsule; LHA, lateral hypothalamic area; VMH, ventromedial hypothalamic nucleus; ZI, zona incerta.

Quantification of Hcrt-immunoreactive neurons revealed no differences between genotypes. Hcrt neuron density was comparable in SNORD116-KO, SNORD115-KO, and SNORD116/115-KO mice relative to their respective WT controls (**Figure 4, B, C and D**).

In contrast, density of MCH-expressing neurons was significantly reduced in SNORD116-KO mice (~12.5% compared to WT). A similar reduction (~16.5%) was observed in SNORD116/115 double-KO mice, whereas MCH neuron density remained unchanged in SNORD115-KO mice. This reduction did not parallel the sleep phenotype: increased REM sleep was observed only in SNORD116-KO mice but not in the double SNORD-KO model, despite a comparable decrease in MCH neuron density (**Figure 4, E, F, G and H**). These results indicate that changes in MCH neuron density alone are not sufficient to account for REM sleep alterations

### Limited changes in hypothalamic gene expression in SNORD-deficient mice at ZT0

Several lines of evidence have suggested a potential link between SNORD116 loss and circadian dysfunction, including the identification of a large number of differentially expressed genes (DEGs) in the hypothalamus of SNORD116-KO mice at ZT0 (24,55,58). This prompted us to examine the impact of SNORD116 and/or SNORD115 deficiency on gene expression in hypothalamic tissues collected at this circadian time point. As shown in **Figure 5A**, none of the three SNORD-deficient models exhibited widespread alterations based on mRNA-seq analysis. Consistent with previous reports, Ndn was mis-expressed upon SNORD116 loss (26,59,60). In addition, changes in Nup210 expression were detected in SNORD115-KO mice, in agreement with previous observations at ZT6 (61). Pomc expression was reduced in SNORD116-KO mice but not in SNORD116/115 double-KO mice, whereas Npy expression was increased specifically in SNORD116/115-KO mice. Overall, these results contrast with the extensive alterations reported by Pace et al. (**Figure 5B**), where several key markers of hypothalamic identity and function exhibit large fold changes and are nearly absent in KO samples (**Figure 5C**). Altogether, our findings indicate that combined deletion of SNORD115 and SNORD116 did not markedly enhance the effects observed upon SNORD116 loss alone.

**Figure 5.**
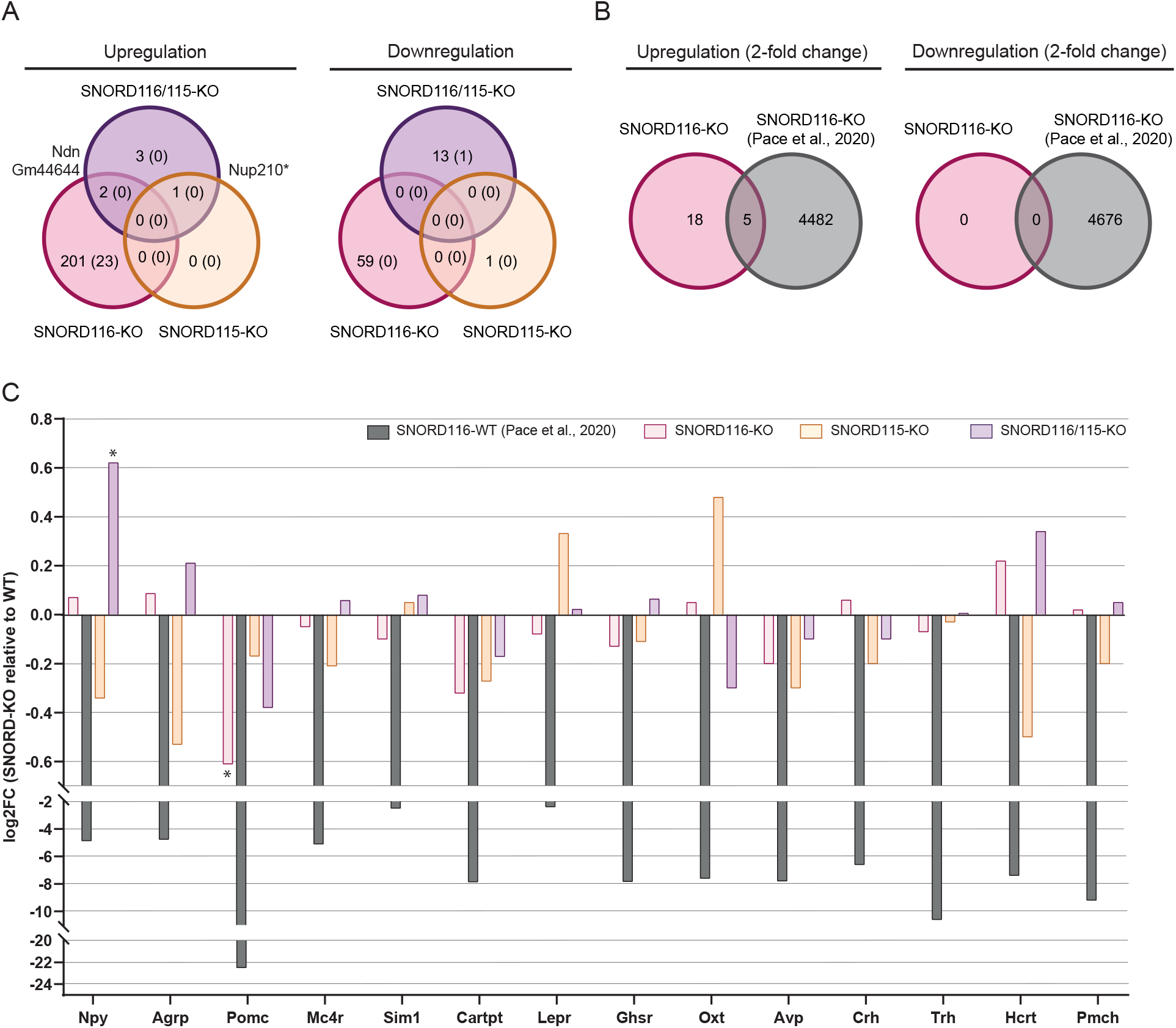
SNORD loss is associated with limited changes in hypothalamic gene expression at ZT0. (**A**) Venn diagram comparing DEGs identified at ZT0 in the hypothalamus of SNORD116-KO, SNORD115-KO, and SNORD116/115-KO mice. The number of DEGs with a fold change >2 is indicated in brackets. (*) *Nup210* falls just below the statistical threshold (adjusted p = 0.055). A small number of poorly annotated transcripts originating from the deleted region (or adjacent regions), which were downregulated in KO samples, were excluded from the DEG counts. Gm44644 is a poorly annotated transcript located outside the deleted interval, mapping between Snrpn and Ndn. Note that *Magel2 et Ube3A* was found to be slightly downregulated (log_2_FC = −0.26) and upregulated, respectively, in SNORD116-KO mice (log2FC = 0.26), but not in SNORD116/115-KO mice. Venn diagrams showing the limited overlap between DEGs identified in SNORD116-KO mice from this study and those reported by Pace et al. with only five common genes (*Shox2, Epha1, Rasl11a, Cdh3*, and *Ramp3*) (**C**) Bar plots showing changes in mRNA levels (log2-fold change, KO relative to WT) of selected markers of hypothalamic identity and functional regulators across SNORD115-KO, SNORD116-KO, and SNORD116/115 double-KO mouse lines. (*) *Npy* is upregulated (log_2_FC = 0.6; padj 7.9e-5) in SNORD116/115-KO mice while *Pomc* is downregulated (log_2_FC = −0.6; padj = 0.02) in SNORD116-KO.

## Discussion

In this study, we found that SNORD116 deletion selectively affected REM sleep without altering SWS architecture or delta power, arguing against a global increase in sleep drive, whereas SNORD115-KO mice did not exhibit detectable sleep phenotypes. Unexpectedly, combined deletion of SNORD116 and SNORD115 genes did not exacerbate the REM sleep phenotype observed in SNORD116-KO, but instead abolished it, as sleep alterations were no longer evident under either baseline conditions or following sleep deprivation. Thus, the effects of SNORD115 and SNORD116 deletions are not additive, arguing against the functional redundancy initially hypothesized, and instead point to a dependency of the SNORD116 phenotype on genetic context.

A similar lack of correspondence emerges at the neuroanatomical level. We observed a reduction in MCH neuron density in both SNORD116-KO and SNORD116/115-KO mice (12.5% and 16.5%, respectively) whereas Hcrt neurons were preserved across all genotypes. Given the central role of the Hcrt system in arousal regulation and its implication in hypersomnia (40–43), the absence of alterations in Hcrt neuron density further supports the dissociation between hypothalamic neuronal population and sleep phenotype. Importantly, this structural alteration did not track the behavioral phenotype: REM sleep changes were present only in SNORD116-KO mice, but absent in double-KO mice despite a comparable reduction in MCH neuron density in the LHA. This dissociation indicates that a decrease in MCH neuron number alone is not sufficient to predict REM sleep alterations, which appears inconsistent with the established role of MCH neurons in promoting REM sleep through inhibition of wake-promoting circuits (46,47,49,50).

Beyond their role in REM sleep regulation, MCH neurons have been implicated in feeding behavior, motivational processes, and aspects of cognitive regulation (62–64). This is particularly relevant in the context of PWS, which is characterized by hyperphagia as well as cognitive and behavioral alterations (1). In this framework, the reduction in MCH neuron density observed in both SNORD116-KO and SNORD116/115-KO mice may reflect alterations extending beyond sleep to other behavioral domains. Notably, this reduction has not consistently been reported across studies, with some reports instead describing a reduction in Hcrt neurons (55). Together, these observations indicate that the functional significance of MCH alterations remains unresolved.

Interpretation of adult phenotypes in SNORD116-deficient models should take developmental context into account. Indeed, loss of SNORD116 - but not SNORD115 - leads to growth, metabolic and hormonal impairments at birth that are associated with variable perinatal mortality, ranging from 0-15% up to 75% in others (22,23,65). This implies that adult analyses are performed on a selected subset of survivors. Given that early-life challenges can shape long-term physiological and behavioral outcomes (66), this selection bias may influence the severity of adult phenotypes and limit the ability to detect primary alterations in sleep regulation and hypothalamic neuronal populations.

At the molecular level, our gene expression analyses at ZT0 identified only a limited number of differentially expressed genes across all SNORD-deficient models, in contrast with previous findings reported at the same circadian time point in SNORD116-KO mice (55). We do not have a clear technical or biological explanation for this pronounced discrepancy. However, our results are consistent with several independent studies reporting limited effects following SNORD116 loss across multiple brain contexts, including adult hypothalamus (59), perinatal hypothalamus (23), laser-captured hypothalamic nuclei (26), and whole brain (60). We therefore favor a parsimonious interpretation in which the differences between studies primarily reflect variations in experimental or environmental conditions, rather than a reproducible transcriptome signature of SNORD116 deficiency. Regardless of the origin of these discrepancies, combined loss of SNORD115 and SNORD116 did not markedly exacerbate the phenotype observed upon SNORD116 deletion alone, further arguing against functional redundancy between these two SNORD. This apparent dissociation between limited mRNA-level changes and measurable anatomical and behavioral alterations suggests that SNORD function may rely on subtle, spatially restricted, or developmental mechanisms that are not easily captured by bulk analyses of adult tissue.

SNORD115 deletion alone has no detectable impact on sleep architecture or homeostatic regulation in young adult mice. A modest increase in SWS episode average duration was, however, observed in older animals, suggesting a possible age-dependent effect, but this effect was limited and did not extend to REM sleep. To date, SNORD115 deficiency has not been associated with overt growth, metabolic, or behavioral phenotypes (61,67). It therefore remains unclear how deletion of SNORD115 results in an apparent normalization of sleep alterations in SNORD116-KO mice. While elucidating this paradoxical genetic interaction is beyond the scope of the present study, several conceptual frameworks can be considered. One possibility is that developmental compensation activated in the double-KO restores sleep regulatory balance, masking the effect of SNORD116 deficiency. Alternatively, SNORD115 and SNORD116 may regulate partially overlapping or functionally opposing pathways, such that removal of SNORD115 mitigates the imbalance induced by SNORD116 loss. Extending this idea, one may hypothesize that a key function of SNORD116 is to constrain SNORD115 activity within neural circuits regulating sleep. In this scenario, the sleep abnormalities observed in SNORD116-KO would arise not primarily from loss of SNORD116 function *per se*, but also from a secondary imbalance driven by SNORD115. Although speculative, this framework provides a coherent interpretation of the interaction observed here.

In conclusion, despite compelling evidence implicating the loss of SNORD116 in the etiology of PWS (4–10,22,23,54–56), our findings highlight a broader difficulty: the apparent absence or inconsistency of phenotypes in SNORD-deficient mouse models may not reflect a lack of biological relevance, but rather the limitations of the experimental and conceptual frameworks used to interrogate them. While REM sleep alterations can be observed in specific genetic contexts, they are subtle, not consistently observed across models, and do not recapitulate key clinical features such as hypersomnolence or hypocretin-related dysfunction. More broadly, our study also underscores the need to investigate mouse models exhibiting simultaneous loss of multiple genes, as recently done for Ndn and Magel2 (68). From a translational perspective, our findings call for a cautious interpretation of current SNORD-deficient mouse models when studying sleep abnormalities in PWS, particularly given that both SNORD116 and SNORD115 are absent in the vast majority of patients.

## Methods

### Animals

All animal procedures were approved by the Institutional Animal Care and Use Committee of the University of Toulouse and the Ministry of Higher Education and Research (APAFIS #36977-202204151214375 v5). SNORD115 and SNORD116/115 knockout (KO) mouse lines were generated in our laboratory using CRISPR-Cas9 technology (61) (Marty et al, manuscript in preparation, see also **Supplementary Material**). SNORD116-KO mouse line was provided by the Institute of Experimental Pathology/Molecular Neurobiology (Münster, Germany) (22). Experimental animals were generated by crossing WT C57BL/6J females with heterozygous SNORD-KO males. Genotypes were determined by PCR analysis of ear biopsies. WT littermates were used as control. All lines were backcrossed for at least twelve generations onto a C57BL/6J background. Only male mice were used. Mice were housed under a 12:12 h light-dark cycle (lights on at 7:30 a.m.; ZT0) at 23 ± 1 °C with *ad libitum* access to food and water.

### Polysomnographic recording

Sleep recordings were performed in 2–3-month-old male mice from the SNORD116, SNORD115, and SNORD116/115 lines (WT and KO, n = 8 per group). Additional recordings were obtained from older cohorts (**Supplemental Figure S4**), including SNORD115 mice aged 7–8 months (WT: n = 10; KO: n = 12) and SNORD116/115 mice aged 6 months (WT: n = 6; KO: n = 6) (**Supplemental Figure S3**).

Animals were habituated to the recording environment for one day prior to data acquisition. EEG/EMG recordings were then collected for 24 h under baseline conditions. To assess sleep homeostasis, a 4-hour sleep deprivation (SD) protocol was applied from ZT0 to ZT4 using gentle handling (including light cage tapping and tactile stimulation with a soft brush). SD was performed in all animals from the 2-3-month-old cohorts. In older cohorts, SD was performed only in SNORD115 mice (WT: n = 6; KO: n = 9), whereas SNORD116/115 mice (WT n = 6; KO n = 6) were recorded under baseline conditions only. Following SD, mice were allowed to sleep *ad libitum*, and recordings continued for an additional 20 h to monitor rebound sleep. Signals were acquired using Spike2 software (version 7.11, Cambridge Electronic Design, UK). Vigilance states (wake, SWS, and REM sleep) were scored in 5-sec epochs according to established criteria (69).

### Spectral Analysis

EEG spectral analysis was performed using Fast Fourier Transform (FFT). Signals were analyzed in the 0.5-50 Hz frequency range with a sampling rate of 100 Hz, in accordance with the Nyquist criterion. FFT was computed on 3-s epochs extracted from scored vigilance states. Spectral analyses were restricted to recordings with stable signal quality; therefore, the number of animals included in these analyses is lower than in the full polysomnographic dataset due to signal quality-based exclusion criteria.

Power spectra were calculated separately for SWS and REM sleep using 2-h time bins. For SNORD115 (WT: n = 5; KO: n = 3) and SNORD116/115 mice (WT: n = 4; KO: n = 4), analyses focused on the ZT4.5-ZT6.5 window under both baseline conditions and post-SD conditions. To isolate rebound sleep, spectral analysis began 30 min after the end of SD.

For each vigilance state, spectral power density was computed across the 0.5–50 Hz frequency range. Delta (0.5–4 Hz) power during SWS and theta (4–12 Hz) power during REM sleep were quantified as the area under the curve of the power spectrum within the respective frequency bands. These values were then normalized to the total spectral power (0.5–50 Hz) of the corresponding vigilance state for each animal. Analyses were performed using a custom MATLAB script (FunSy, LNCA, Strasbourg).

### Quantification of Hcrt and MCH neurons

Hcrt and MCH neurons were quantified in male mice aged 4-6 months from SNORD115 (WT, n = 6; KO, n = 6), SNORD116 (WT, n = 5; KO, n = 5), and SNORD116/115 (WT, n = 7; KO, n = 6) lines. Coronal sections spanning −1.06 mm to −1.94 mm from bregma were analyzed according to the Paxinos atlas, with 250 µm spacing between sections. Three sections per animal were used for Hcrt quantification and four for MCH quantification.

Images were acquired at 10x magnification (Leica DM6000B). Regions corresponding to the LHA were delineated using Qupath software. Immunopositive neurons were manually counted blind to genotype and expressed as cell density relative to the estimated volume of the analyzed LHA. No differences in LHA volume were observed across genotypes (see **Supplemental Figure S6**).

### Transcriptomic analyses

Animals were rapidly euthanized by cervical dislocation at ZT0, and hypothalamic tissues were immediately harvested, snap-frozen in liquid nitrogen, and stored at −80°C. Total RNA was extracted using TRI Reagent (Euromedex) and treated with RNase-free RQ1 DNase (Promega) and Proteinase K (Sigma) to remove DNA and proteins. Library preparation (polyA+ selection) and Illumina sequencing (2 × 150 bp configuration) were performed by Genewiz. Raw sequencing reads (fastq) were quality-checked using FastQC. Reads were aligned to the reference mouse genome (GRCm39) using STAR aligner (STAR_2.5.4a), and gene-level counts obtained based on GENCODE annotations (GRCm39). Count matrices were filtered using the Bioconductor HTSfilter (s.min = 1, s.max = 50) and differential expression analysis (including normalization and calculation of log2 fold changes and adjusted p-values) was performed using Bioconductor DESeq2 (v1.42.0), with each mutant genotype (KO) compared independently to WT controls. Raw data generated in this study are available in the Sequence Read Archive (SRA) database under accession number XXXXXXXX.

### Statistical analysis

Statistical analyses were performed using GraphPad Prism 9. Data distribution was assessed using the Shapiro-Wilk test. Vigilance state distributions, episode counts, and sleep parameters across time bins and light/dark phases were analyzed using two-way repeated-measures ANOVA or Mixed-effects model (REML) (Genotype x Time or Genotype x Phase) with Geisser-Greenhouse’s correction followed by Šídák’s post hoc tests when appropriate. Spectral data and other pairwise comparisons were analyzed using unpaired t-tests with Welch’s correction or Mann-Whitney tests depending on data distribution.

For Hcrt and MCH cell density, group comparisons were performed using unpaired t-tests or Mann-Whitney tests as appropriate.

Data are presented as mean ± SEM. Statistical significance was set as P < 0.05.

## Author Contributions

JCR, JC and LV designed the study. JCR and VM performed the experiments. JCR, JC and LV analyzed the data. JCR, VM, JC and LV interpreted the results. JCR, JC and LV wrote the manuscript. JC and LV supervised the project. All authors approved the final version of the manuscript.

## Acknowledgments

Mice were housed in the ABC Facility of ANEXPLO, Toulouse. The authors thank Camille Lejards, Anna B. Szabo, Lionel Dahan, Patrick Arrufat, Valentine Guidolin, Orane Azulay-Fournier, and Samir Boutaleb for technical help. The script used for spectral analysis was kindly provided by Romain Goutagny (Funsy, LNCA, Strasbourg), and edited with the help of Alid Al-Asmar and Tom Orjollet--Lacomme. This work was supported by the Centre National de la Recherche Scientifique (CNRS), the University of Toulouse, the ANR (ANR-22-CE12-0020 to J.C.). J.C.R. received a Ph.D. fellowship from the French Ministry of Research.

