## Supplemental for "Loss of SNORD115 mitigates SNORD116-driven sleep abnormalities in mouse models of Prader-Willi syndrome"

This PDF includes:

***Figure S1. SWS and REM episode duration are not altered by genotype under baseline conditions.***
***Figure S2. SWS and REM episode duration are not altered by genotype following sleep deprivation.***
***Figure S3. Sleep architecture of SNORD116/115 mice at 5 months under baseline conditions.***
***Figure S4. Sleep architecture under baseline conditions and following 4-hour sleep deprivation in SNORD115-KO mice.***
***Figure S5. Absence of SNORD115 and SNORD116 expression in Hcrt and MCH neurons of male SNORD116/115-KO mice.***
***Figure S6. Total lateral hypothalamic area (LHA) volume across genotypes.***

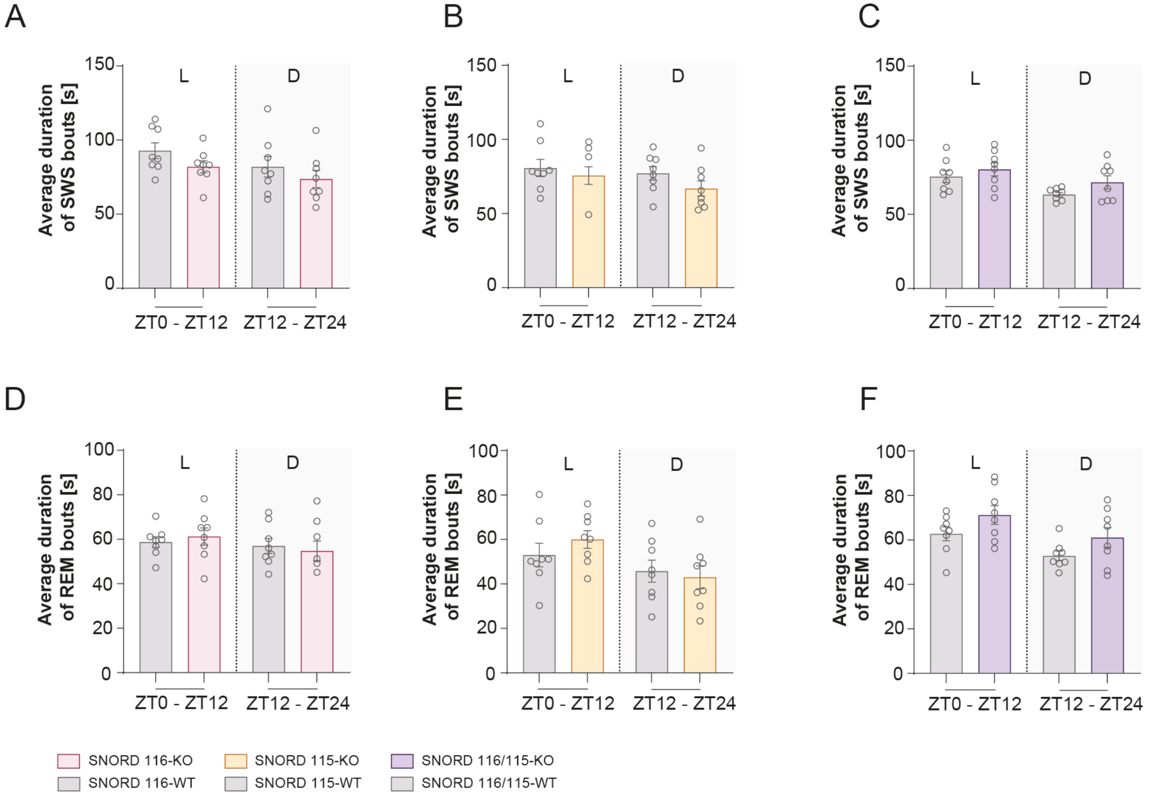
A
B
C
D
E
F
Average duration of SWS bouts [s]
Average duration of REM bouts [s]
L
D
ZT0 - ZT12
ZT12 - ZT24
SNORD 116-KO
SNORD 116-WT
SNORD 115-KO
SNORD 115-WT
SNORD 116/115-KO
SNORD 116/115-WT

***Figure S1. SWS and REM episode duration are not altered by genotype under baseline conditions.*** *Sleep episode duration was assessed under baseline conditions in SNORD116-KO, SNORD115-KO, and SNORD116/115 double-KO mice and their WT littermates. (**A–C**) Mean slow-wave sleep (SWS) episode duration (sec) across light (L) and dark (D) phases in SNORD116-KO (**A**), SNORD115-KO (**B**), and SNORD116/115 -KO (**C**) mice. (**D–F**) Mean REM sleep episode duration (sec) across light and dark phases in SNORD116-KO (**D**), SNORD115-KO (**E**), and SNORD116/115-KO (**F**) mice. Data are presented as mean ± SEM. Statistical analyses were performed using two-way ANOVA (Genotype x Phase).*

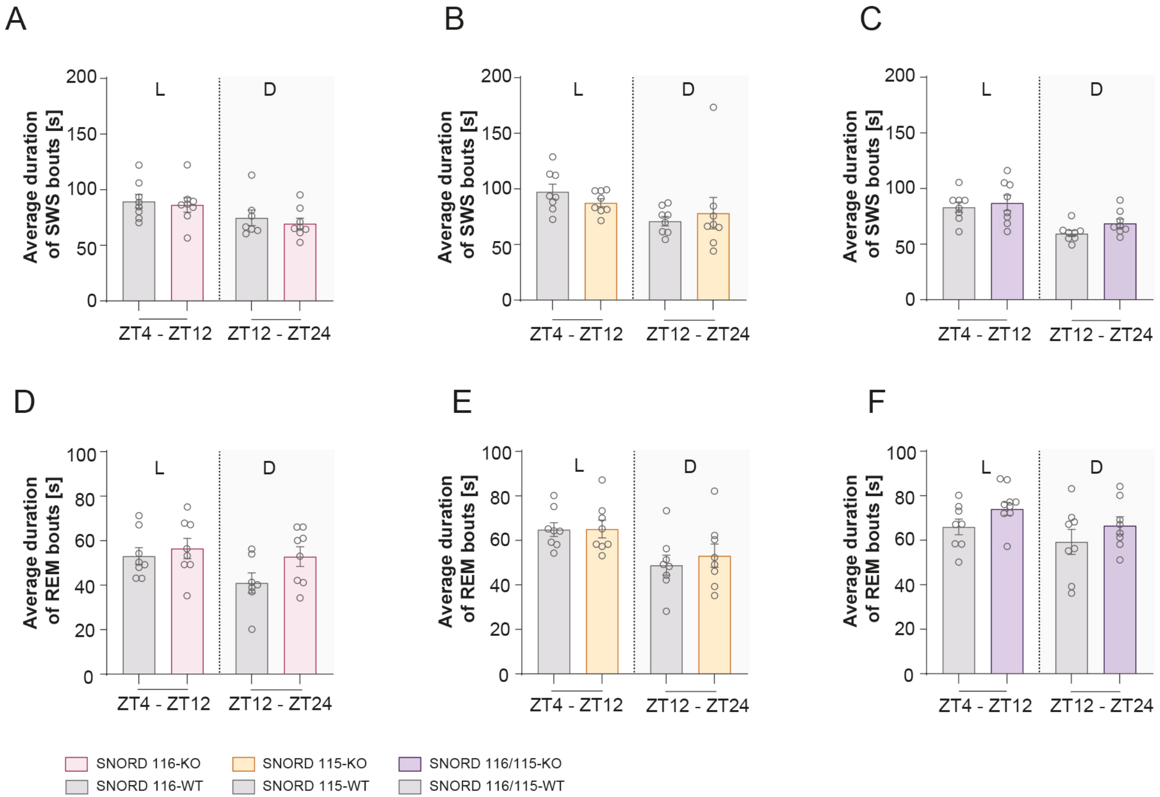
A
B
C
D
E
F
Average duration of SWS bouts [s]
Average duration of REM bouts [s]
L
D
ZT4 - ZT12
ZT12 - ZT24
SNORD 116-KO
SNORD 116-WT
SNORD 115-KO
SNORD 115-WT
SNORD 116/115-KO
SNORD 116/115-WT

***Figure S2. SWS and REM episode duration are not altered by genotype following sleep deprivation.*** *Sleep episode duration were assessed during the rebound period following sleep deprivation in SNORD116-KO, SNORD115-KO, and SNORD116/115 double-KO mice and their WT littermates. (**A–C**) Mean slow-wave sleep (SWS) episode duration (sec) across light (L) and dark (D) phases in SNORD116-KO (**A**), SNORD115-KO (**B**), and SNORD116/115-KO (**C**) mice. (**D–F**) Mean REM sleep episode duration (sec) across light and dark phases in SNORD116-KO (**D**), SNORD115-KO (**E**), and SNORD116/115-KO (**F**) mice. Data are presented as mean ± SEM. Statistical analyses were performed using two-way ANOVA or mixed-effects models (Genotype x Phase*).

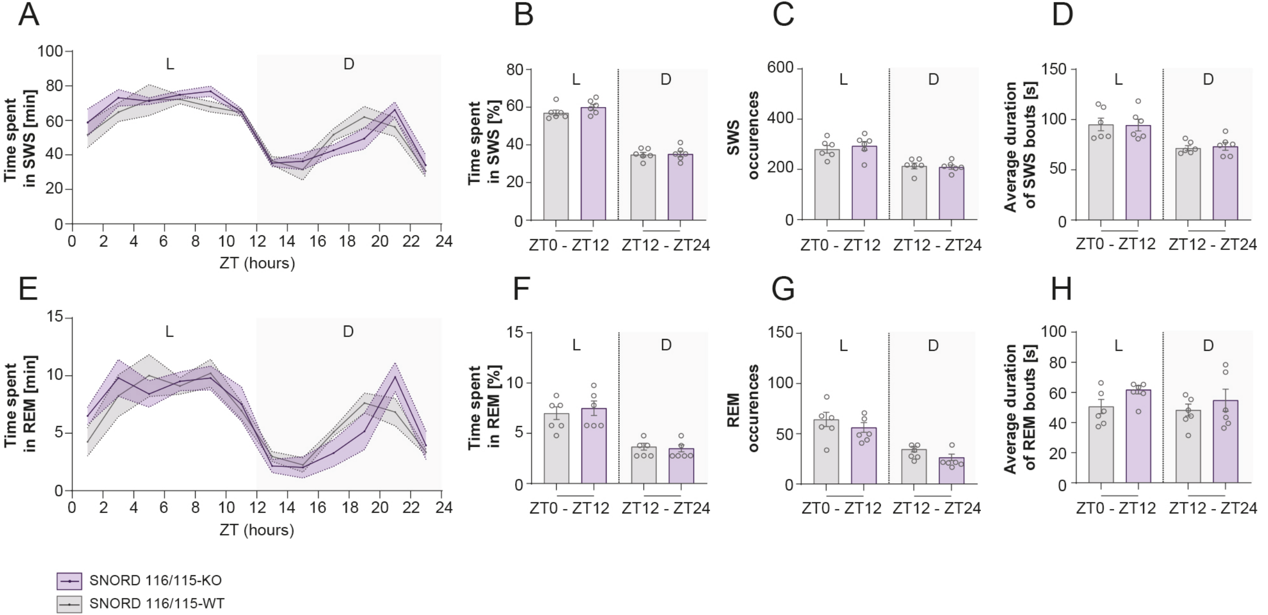

A
B
C
D
E
F
G
H
Time spent in SWS [min]
Time spent in SWS [%]
SWS occurences
Average duration of SWS bouts [s]
Time spent in REM [min]
Time spent in REM [%]
REM occurences
Average duration of REM bouts [s]
ZT (hours)
L
D
ZT0 - ZT12
ZT12 - ZT24
SNORD 116/115-KO
SNORD 116/115-WT

***Figure S3. Sleep architecture of SNORD116/115 mice at 5 months under baseline conditions.*** *Polysomnographic recordings were performed in 5-month-old SNORD116/115 double-KO mice and their WT littermates under baseline conditions. (**A–D**) Slow-wave sleep (SWS) parameters. (**A**) Time spent in SWS (min) across the 24-hour cycle (ZT0–ZT24). (**B**) Percentage of time spent in SWS across light (L) and dark (D) phases. (**C**) Number of SWS episodes. (**D**) Mean SWS episode duration. (**E–H**) Rapid eye movement (REM) sleep parameters. (**E**) Time spent in REM sleep across ZT bins. (**F**) Percentage of REM sleep across L/D phases. (**G**) Number of REM episodes. (**H**) Mean REM episode duration. Data are presented as mean ± SEM. Statistical analyses were performed using two-way ANOVA (Genotype x Zeitgeber Time [ZT] or Genotype x Light/Dark phase [L/D]).*

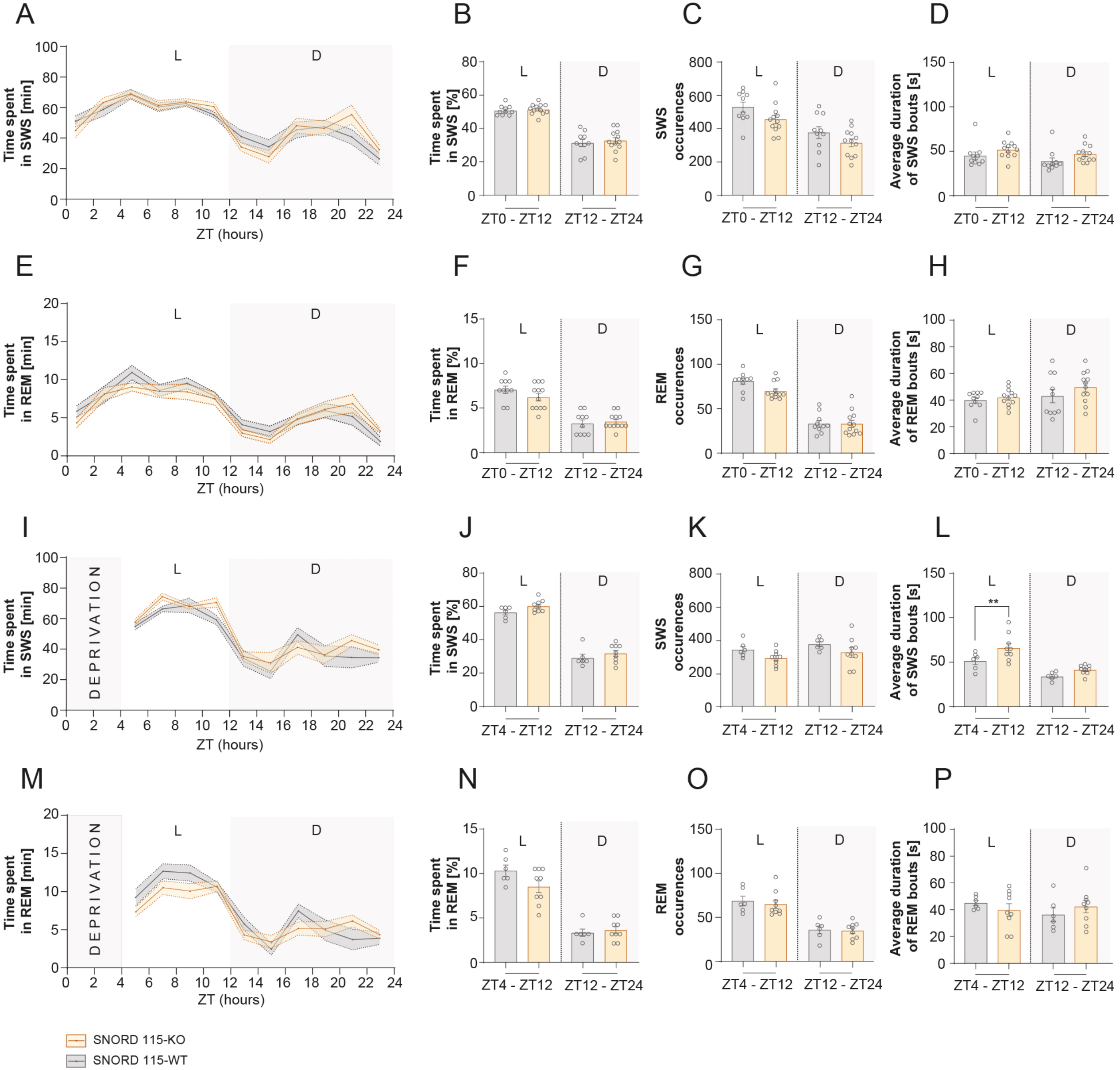
A
B
C
D
E
F
G
H
I
J
K
L
M
N
O
P
L
D
Time spent in SWS [min]
Time spent in REM [min]
Time spent in SWS [%]
Time spent in REM [%]
SWS occurences
REM occurences
Average duration of SWS bouts [s]
Average duration of REM bouts [s]
ZT (hours)
ZT0 - ZT12
ZT12 - ZT24
ZT4 - ZT12
DEPRIVATION
**
SNORD 115-KO
SNORD 115-WT

***Figure S4. Sleep architecture under baseline conditions and following 4-hour sleep deprivation in SNORD115-KO mice.*** *Polysomnographic recordings were performed in 8-month-old SNORD115-KO mice and their WT littermates under baseline conditions and following 4-hour sleep deprivation (SD). (**A–D**) Baseline SWS parameters. (**A**) Time spent in slow-wave sleep (SWS; min) across the 24-hour cycle (ZT0–ZT24). (**B**) Percentage of time spent in vigilance states across light and dark phases. (**C**) Number of SWS episodes. (**D**) Mean SWS episode duration. (**E–H**) Baseline REM sleep parameters. (**E**) Time spent in REM sleep across ZT bins. (**F**) Percentage of REM sleep across light and dark phases. (**G**) Number of REM episodes. (**H**) Mean REM episode duration. (**I–L**) SWS homeostasis following sleep deprivation. Same parameters as in baseline (**A–D**). (**M–P**) REM sleep homeostasis following sleep deprivation. Same REM parameters as in E–H. Data are presented as mean ± SEM. Statistical analyses were performed using two-way ANOVA (Genotype x Zeitgeber Time [ZT] or Genotype x Light/Dark phase [L/D]), followed by Šídák's post hoc tests when appropriate. **P < 0.05; **P < 0.01.*

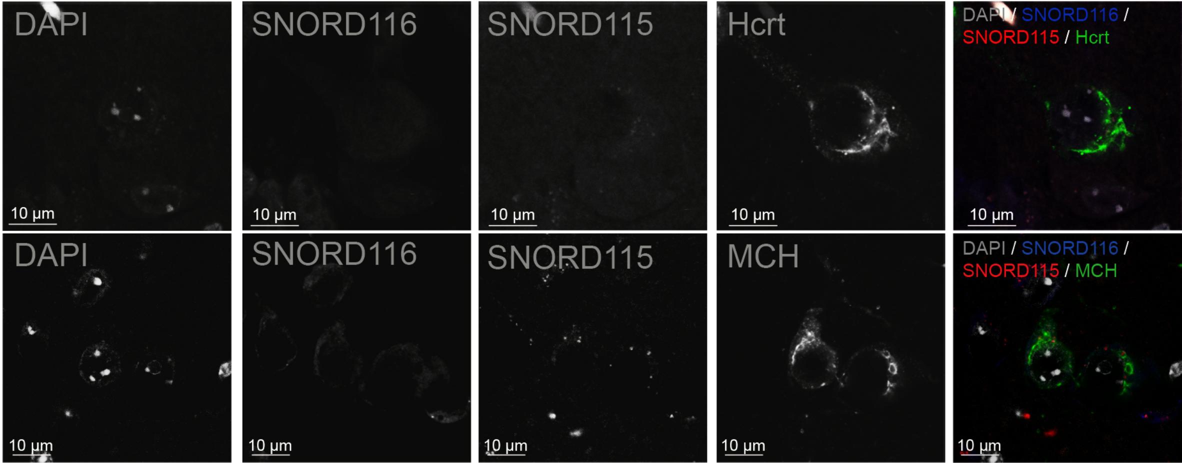
DAPI
SNORD116
SNORD115
Hcrt
DAPI / SNORD116 / SNORD115 / Hcrt
MCH
DAPI / SNORD116 / SNORD115 / MCH
10 µm

***Figure S5. Absence of SNORD115 and SNORD116 expression in Hcrt and MCH neurons of male SNORD116/115-KO mice***. *A relatively high background signal was observed in both the nucleoplasm and cytoplasm, particularly with the SNORD116 probe. These signals should not be considered specific, as they were also present in brain sections from SNORD116/115-KO mice used as negative controls. In contrast, the nucleolar signals detected with anti-SNORD115 and anti-SNORD116 probes in wild-type samples were absent in KO sections, confirming the specificity of the probes.*

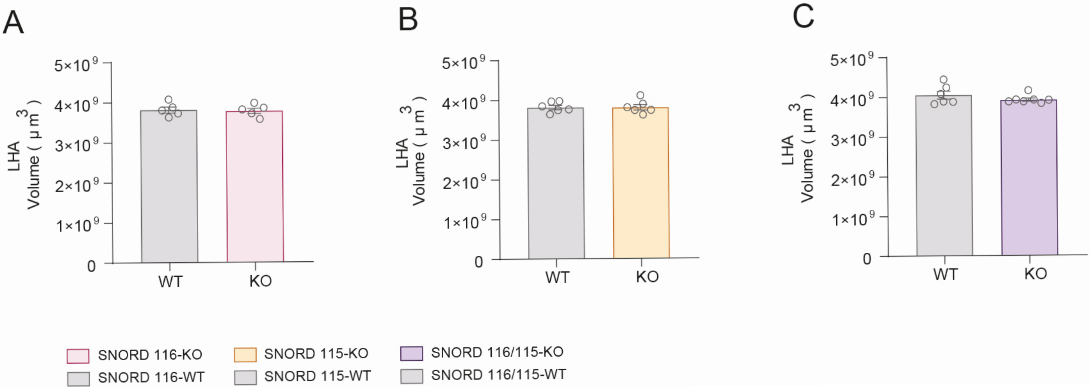
A
B
C
LHA
Volume ( μ m 3 )
5×10 9
4×10 9
3×10 9
2×10 9
1×10 9
0
WT
KO
SNORD 116-KO
SNORD 116-WT
SNORD 115-KO
SNORD 115-WT
SNORD 116/115-KO
SNORD 116/115-WT

***Figure S6. Total lateral hypothalamic area (LHA) volume across genotypes.*** *Total LHA volume (µm³) was quantified in SNORD116 (**A**), SNORD115 (**B**), and SNORD116/115 mouse (**C**) lines. Statistical analyses were performed using unpaired t-tests or Mann–Whitney tests, depending on data distribution. Data are presented as mean ± SEM.*

**Supplementary materials for**

### Loss of SNORD115 mitigates SNORD116-driven sleep abnormalities in mouse models of Prader-Willi syndrome

Julie C. Riviere[1,2,*], Virginie Marty[2], Jérôme Cavaillé[2,*], and Laure Verret[1,*]

[1] Univ Toulouse, CNRS, Centre de Recherches sur la Cognition Animale, Centre de Biologie Intégrative, Toulouse, France

[2] Univ Toulouse, CNRS, Molecular Cellular Development, Centre de Biologie Intégrative, Toulouse, France

This PDF includes:

Generation of SNORD116/115-KO and SNORD115-KO.
Electrode Preparation
Animal surgery for EEG
Polysomnographic set up
Tissue preparation
Immunohistochemistry (MCH and Hcrt)
Fluorescent *in situ* hybridization (FISH) combined with immunofluorescence

#### Methods

##### Generation of SNORD116/115-KO and SNORD115-KO.

The double KO mouse model lacking both SNORD116 and SNORD115 was generated using the CRISPR-Cas9 strategy previously described for the SNORD115-KO line (61). Cas9 mRNA (GeneArt CRISPR nuclease mRNA, Invitrogen) and two guide RNAs (MEGAshortscript T7 Transcription Kit; Ambion) designed to base-pair upstream and downstream of SNORD116-Ipw-SNORD115 gene array (crispr.mit.edu tool) were co-injected into zygotes obtained from superovulated (C57BL/6J × CBA) F1 females mated with F1 males. A few picoliters of Cas9 mRNA (100 ng/ml) and sgRNAs (25 ng/ml each) were co-injected into the cytoplasm and pronuclei of zygotes. Injected embryos were cultured in M16 medium until the blastocyst stage to assess cleavage efficiency or were directly transferred into the oviductal ampullae of pseudopregnant recipient females. The full description of the perinatal and adult phenotype will be reported elsewhere (Marty et al, manuscript in prep).

##### Electrode Preparation

EEG electrodes consisted of Teflon-coated silver wire, while reference electrodes were made of silver. EMG electrodes consisted of multi-stranded stainless-steel wire coated with perfluoroalkoxy, with the distal end inserted into neck muscles. All electrodes were soldered to miniature gold-plated pins and secured onto a six-channel plastic connector (PlasticsOne) using methyl methacrylate-based dental cement (Duralay). Electrode integrity and conductivity were verified prior to implantation using a multimeter.

##### Animal surgery for EEG

Mice were anesthetized with isoflurane (1000 mg/g/ml; Vetflurane, Virbac) and placed in a stereotaxic frame (Kopf) with continuous gas delivery. Body temperature was maintained at ~36 °C using a heating pad with rectal monitoring. Local anesthesia (lidocaine, 7 mg/kg) was administered prior to a midline scalp incision.

Small burr holes were drilled over the frontal (AP: +1.5 mm; L: -1.5 mm from bregma) and parietal cortices (AP: -2.2 mm; L: -2 mm from bregma), and over the occipital bone for the reference. EEG electrodes were inserted subdurally, and EMG electrodes were placed into the neck muscles. Electrodes were secured using dental adhesive (Superbond, Sun Medical) and dental cement (Duralay).

At the end of surgery, mice received analgesia (Metacam, 2 mg/ml) and were allowed to recover individually under a heat source. A minimum recovery period of 7 days was provided before recordings.

**Polysomnographic set up**

During polysomnographic recordings, mice were housed individually in Plexiglas cages with free access to food and water, and placed inside grounded Faraday cages to minimize electrical noise. Animals were connected via a flexible cable to a commutator (PlasticsOne) through a custom-built voltage follower.

Signals were amplified (AM Systems 3600), referenced, and band-pass filtered (EEG: 0.3–1000 Hz; EMG: 3–20,000 Hz). Data was digitized using a CED Power1401 mk II acquisition system and recorded with Spike2 software. Signals were sampled at 100 Hz.

**Tissue preparation**

Mice were anesthetized with isoflurane (Vetflurane) followed by intraperitoneal injection of sodium pentobarbital (Euthasol, 400 mg/ml). Perfusions were performed between 2:00 p.m and 5:30 p.m to minimize circadian variability. Animals were transcardially perfused with 0.9% saline, and brains were post-fixed in 4% formaldehyde at 4°C. After washing, brains were cryoprotected in 30% sucrose and sectioned coronally (25 µm) using a freezing microtome. Sections were stored in cryoprotectant at −20°C until use. Prior to staining, sections were rinsed in phosphate buffer (0.1M) with 0.9% NaCl (PBS) and treated to reduce endogenous peroxidase activity and autofluorescence (80% PBS, 10% methanol, 3% $H_2O_2$).

**Immunohistochemistry (MCH and Hcrt)**

Sections were incubated in PBS with 0.25% Triton-X (PBST) with normal goat serum (NGS) for anti-MCH immunohistochemistry, whereas no serum was used for anti-Hcrt staining. Sections were incubated overnight with rabbit anti-Hcrt-A (1:10,000; Phoenix Pharmaceuticals, H-003-30) or rabbit anti-MCH antibodies (1:10,000; Phoenix Pharmaceuticals, H-070-47; 5% NGS). After washing, sections were incubated with goat anti-rabbit biotinylated secondary antibodies (1:1,000; Vector Labs, BA1000) and processed using an avidin–biotin complex (ABC, Vector Labs). Signal was revealed using DAB (Vector Labs), and sections were counterstained with cresyl violet.

**Fluorescent *in situ* hybridization (FISH) combined with immunofluorescence**

Sections were processed for immunofluorescence. Blocking was performed in PBST containing 10% normal donkey serum (NDS) for sections incubated with anti-MCH antibodies.

Sections were then incubated overnight at room temperature with rabbit anti-Hcrt-A (1:5,000; Phoenix Pharmaceuticals, H-003-30) or anti-MCH antibodies (1:1,000; Phoenix Pharmaceuticals, H-070-47; 10% NDS). After washing, sections were incubated for 90 minutes with Alexa Fluor 488–conjugated donkey anti-rabbit secondary antibodies (1:500; Invitrogen, A11008).

In situ hybridization was then performed on the same sections. After pre-hybridization (45 min, room temperature) in 2X Saline-Sodium Citrate (SSC) with 15% formamide, sections were hybridized overnight at 37°C in buffer containing 15% formamide, 2X SSC, 10% dextran sulfate, 150 µg/ml yeast tRNA, and Cy3- and Cy5-labeled oligonucleotide probes targeting SNORD115 and SNORD116 (300 ng each). Sections were washed in 15% formamide/2X SSC followed by 1X SSC and mounted in Mowiol containing DAPI (0.1 mg/ml).

Cy3- and Cy5-labelled antisense DNA oligonucleotide probes were purchased from Sigma-Aldrich

SNORD115:CY3-GGGCCTCAGCGTAATCCTATTGAGCATGAATTTTATGTCATCACC-CY3;
SNORD116:CY5-TCAGTTCCGATGAGAGTGGCGGTACAGAGTTTTCACTCATTTTGTTCA-CY5
